# Heterogeneous multi-scale framework for cancer systems models and clinical applications

**DOI:** 10.1101/633933

**Authors:** Alokendra Ghosh, Ravi Radhakrishnan

## Abstract

Clinical Cancer models need to incorporate a wide variety of patient data and tumor heterogeneity which requires integration of multiple models. Due to differences in time and length scales of individual processes, such a model integration is a challenging task. Here we have developed an integrated framework combining ErbB receptor mediated Ras-MAPK and PI3K/AKT pathway with p53 mediated DNA damage response pathway. We have applied this in a clinical setting to predict patient specific response of different treatments in cancers of prostate, lung and kidney.

## I. Introduction

Systems models of key signaling pathways in cancer such as ErbB receptor mediated Ras-MAPK or PI3K/AKT have been extensively used to understand the effect of growth factors, inhibitors, chemotherapeutic agents etc. on tumor cells [1]. These models often consist of a system of nonlinear ordinary differential equations which need to be solved together to predict the effect of mechanical or chemical stimuli (for e.g. drug dosage) in terms of activity of one or more key downstream proteins such as ERK or AKT which are important regulators of cell fate decisions. Although such systems models are greatly useful and have helped uncover important emergent behavior of signaling networks such as ultrasensitivity, bistability and oscillations [2], they miss many key features that would make them useful in a clinical setting. 1) The predictions of activity of proteins such as ERK or AKT cannot be directly translated into a clinically useful cell fate parameter such as cell kill rate. 2) They don’t work as well when there are multiple biological processes operating under different time and length scales such as receptor based signaling (4-6 hours) and cell cycle (24-48 hours). 3) They cannot incorporate important cellular physics like mechanics of the cell membrane, ECM and the cytoskeleton. 4) The parameter space of such models often exhibits sloppy/stiff character (explained in Methods) which affect the accuracy of predictions and the robustness of these models. Here we have developed a multiscale and multiparadigm framework for systems and pharmacodynamic models that helps us address some of the above shortcomings. This framework was used to successfully integrate a single-cell systems model of ErbB receptor mediated Ras-MAPK and PI3K/AKT pathway [1] with a tumor suppressor p53 mediated DNA damage response and cell cycle pathway [3]. The integrated model was used in a clinical setting using gene/protein expression data and drug dosage/schedule information from actual patients of lung and prostate adenocarcinoma. Such multiscale modeling frameworks have great potential in the field of personalized medicine.

## II. METHODS

### A. Description of framework

The heterogeneous multiscale framework consists of one or more systems models of biological processes which are to be run together. The constituent models can have different characteristic time scales and time resolution (discrete or continuous) which determine how the models are to be integrated. The framework also needs a list of species/components which are common to each pair of models which form the interface of the models and mediate the flow of information from one model to the other. If two models do not have any common species, they do not directly interact with each other (the interaction can however come through a third model which shares interfaces with both models). For example, we combined an ErbB receptor mediated Ras-MAPK and PI3K/AKT signaling module and androgen receptor signaling module and a TP53 mediated DNA damage response module in cancers of prostate. Here the ErbB receptor and androgen receptor models have a continuous time description and characteristic time scales of about 6-8 hours. On the other hand, the p53 mediated DNA damage response pathway has a discrete time description (Boolean model) and characteristic time scale of 24-48 hours. Such discrete models are used due to lack of quantitative data on highly complex models like DNA damage response pathways. All the component models have common interfaces with each other which mediate the flow of information, Since the characteristic time scales are well separated we can use pseudo-steady state approximations to combine these models.

### B. Model Interfaces and Hybrid Simulator Algorithm

As mentioned above, when two processes are well-separated in their characteristic time scales, then from the perspective of the slower process the faster process is at steady state [4]. This observation allows us to couple dynamics of these processes by evolving the slower process using steady state information from faster process. When two processes are modeled by different mathematical representations for example continuous time ordinary differential equations and discrete time logical equations, a meaningful coupling of the models can be achieved by identifying a set of species which are common to both and modifying the governing equations (ODE or Boolean rules) and initial state of one model by using information obtained by running the other model for a specified amount of time [5]. The algorithm for the hybrid simulator is shown in the flowchart below and described in [6].

**Fig 1:**
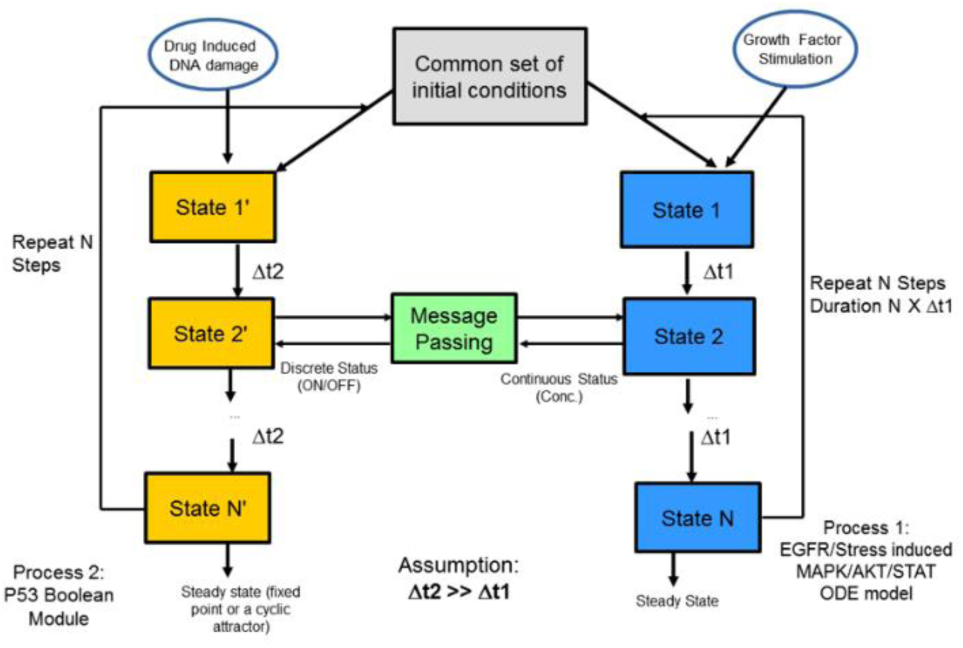
Flowchart showing the algorithm of hybrid simulator. Here we have two different processes (boxes with blue and yellow) with their characteristic time scales Δt1 and Δt2. These are initialized and run for N steps. After each step information is passed between the models to reinitialize them. This is continued until both converge to a common steady state.

### C. Parameter Space Sensitivity, Sloppyness and Robustness

The model parameter space has a profound effect on its predictions. We are interested in how sensitive the predictions are with respect to small perturbations in the parameters. Another related question is how the uncertainties in the parameters contribute to the uncertainties in the model predictions. For complex nonlinear models, finding answers to such questions are nontrivial. We use a variety of different global sensitivity analysis methods like multi parametric sensitivity analysis (MPSA) [7] and determination of model parameter space eigenvalue spectra to characterize sloppiness and robustness of the models [8].

## III. Results

### A. Model Validation

The individual component models of the hybrid multiscale framework were validated individually. However, such individual validation are not sufficient when they are combined and we need to validate the predictions from the overall simulator. To do this we used published [9] molecular and clinical data on prostatectomized patients considered high risk. The model predicted net cell growth rate and production of Prostate Specific Antigen (PSA) was compared with observed values [10]. The model successfully captured qualitative trends like recurrence events in specific patients following prostatectomy.

**Fig 2:**
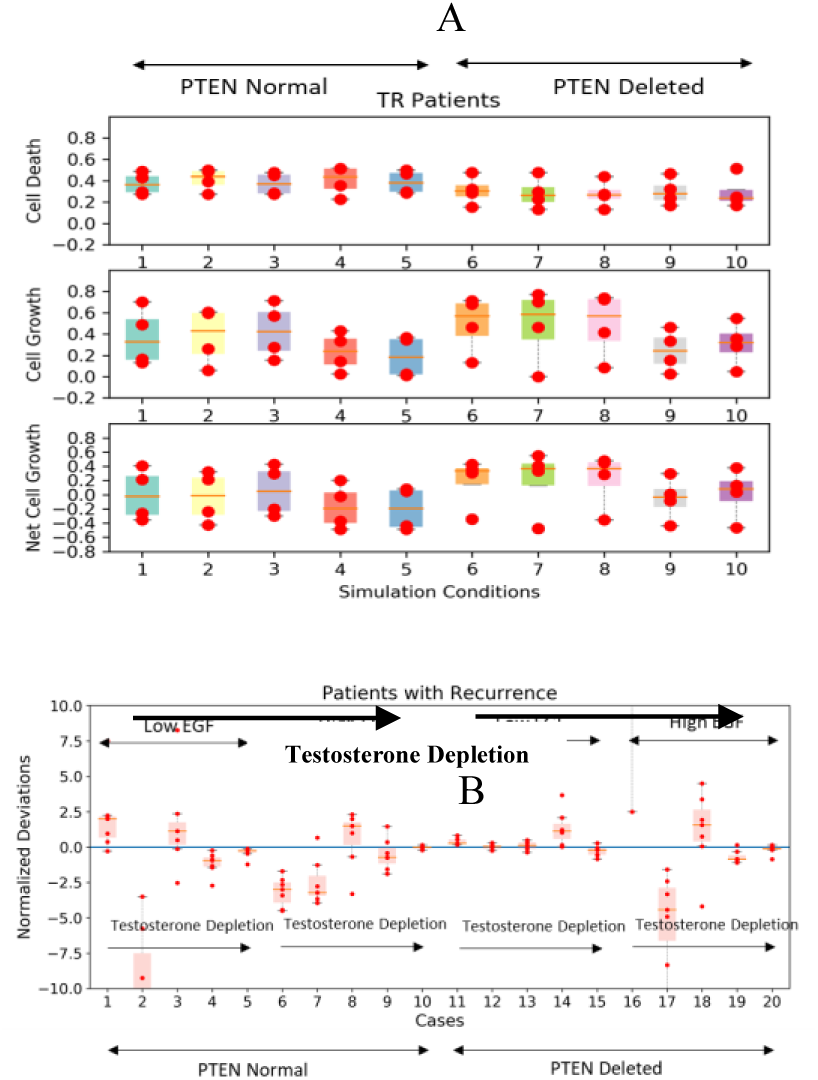
Model predictions of cell kill, growth and net growth probabilities for patients with tumor recurrence (A) and normalized deviation of net cell growth probabilities with respect to control patients for different growth factor and testosterone levels (B)

### B. Effect of PTEN deletion and biochemical recurrence in Prostate Adenocarcinoma

Next, we applied the framework to find answers to some critical questions in prostate adenocarcinoma. Deletion of PTEN, a protein with key regulatory roles in many pathways like PI3K-AKT is often associated with high risk tumors and castration resistant forms of prostate cancer [11]. Tumor recurrence after prostatectomy is often predicted using the serum PSA levels as markers and used to determine whether adjuvant therapies like Androgen-Deprivation Therapy (ADT) need to be administered or not. To determine the effect of PTEN deletion and whether PSA levels are truly significant or not we analyzed data on prostate cancer patients from TCGA. A subset of the patients with high Gleason Scores and those who did not undergo any neoadjuvant therapies or radiation therapies were selected. We classified the patients into groups of control, patients with biochemical recurrence (indicated by PSA values) and patients with tumor recurrence. We used differential gene expression analysis to determine which genes are over/under expressed in our networks and obtained predicted net cell growth for these patients. One of the key observations was that the tumor recurrence events were not simply dependent on single markers like PTEN deletion but depends on differences in expressions of multiple genes/proteins, signaling network dynamics and tumor microenvironment heterogeneity. In particular variabilities in tumor microenvironment was seen to play a key role in determining the effect of ADT [10].

## Acknowledgment

We thank the CHIC consortium members for inputs and helpful discussions.

